# The insect- and plant-associated lifestyles of *Pseudomonas protegens* CHA0 are preserved following serial passage through insect larvae

**DOI:** 10.64898/2026.03.19.712869

**Authors:** Maria Zwyssig, Jana Schneider, Gijs Selten, Christoph Keel, Monika Maurhofer, Ronnie de Jonge

## Abstract

The plant-beneficial bacterium *Pseudomonas protegens* CHA0 (CHA0) is widely studied for the biological control of soil-borne plant diseases. Beyond its root-colonising capabilities, CHA0 can also infect and kill insect larvae and thus exhibits a multi-host lifestyle shared with other plant- and insect-colonising bacteria. To better understand the robustness of this multi-host lifestyle, we subjected CHA0 to ten consecutive passages through larvae of the pest insect *Plutella xylostella* via repeated cycles of insect colonisation and killing forcing it into an insect-only lifestyle. Overall, serial passaging did not result in consistent changes in insect killing speed, larval or root colonisation, plant protection efficiency, microbial antagonism or *in vitro* growth. This suggests that its multi-host lifestyle was conserved following serial passage. Nonetheless, a few independently passaged lines showed an increase in larval killing speed, which in one case might be linked to choline uptake. To disentangle changes specific to the insect host from those arising due to the experimental system itself, we conducted parallel serial passages through the same system while omitting the insect host. In some of these lines, exposure to the background of the system led to changes in microbial antagonism and in *in vitro* growth, which likely are associated with mutations in regions encoding for regulatory systems. Our findings indicate that *P. protegens* CHA0 remains phenotypically stable in complex environments such as an insect host, suggesting that the multi-host lifestyle might also be conserved when applied in the field and supporting CHA0’s potential for reliable biocontrol performance against both plant diseases and insect pests.

**Author summary:** Controlling insect pests with living organisms, known as biological control, offers an environmentally friendly alternative to chemical pesticides. The plant-beneficial bacterium *Pseudomonas protegens* CHA0 is a promising biocontrol candidate that not only colonizes plant roots but also infects and kills certain insect larvae. This ability to colonize different hosts appears to be a conserved trait also observed in other bacteria. To better understand the robustness of this multi-host lifestyle, we repeatedly exposed CHA0 to larvae of the insect pest *Plutella xylostella* and assessed the resulting physiological and genetic changes. Surprisingly, after ten cycles, CHA0 largely retained its insect-killing and plant-protective traits. Although a few populations showed minor changes, including slightly faster insect killing and traits associated with aspects of the experimental system, these changes were limited in scope. Overall, our findings suggest that *P. protegens* CHA0 does not change rapidly in complex environments such as an insect host, supporting its potential for reliable biocontrol performance in the field.

## Introduction

Plant-beneficial members of the genus *Pseudomonas* are of great interest for the biological control of root diseases and insect pests in agriculture [1–3]. They are competitive root-colonising bacteria which can promote plant growth and protect plants against various soil-borne plant diseases [4]. The direct plant beneficial properties of these bacteria are based on plant growth stimulation by the production of phytohormones, e.g., indole acetic acid, nutrient solubilisation, or by increasing plant stress tolerance [5]. Induced systemic resistance, competition for nutrients and space, and direct antibiosis result in the inhibition of plant pathogenic organisms and indirectly contribute to plant health [2,6]. Direct antibiosis of different soil-borne pathogens, for example, is mainly due to the production of antimicrobial agents such as 2-4-diacetylphloroglucinol, pyoluteorin, pyrrolnitrin, phenazines, and hydrogen cyanide [7,8]. Some of the plant-beneficial pseudomonads such as *Pseudomonas protegens* additionally show insecticidal activity against lepidopteran and dipteran insect larvae [9–14], in which lethal infections of the larvae’s haemolymph can occur when *P. protegens* is taken up orally. For this, several obstacles need to be overcome by the bacteria, for which a multifactorial infection mechanism was proposed by Vesga et al. [15]. First, the bacteria need to withstand the harsh gut environment and the resident gut microbiota with the help of different factors such as a type VI secretion system [16], antimicrobial compounds [15,17] or O-antigenic polysaccharides [18].

Through a not yet fully understood mechanism, *P. protegens* is able to breach the gut epithelial barrier of the insect host to access the haemolymph. There, the bacteria secrete the insecticidal Fit toxin, which ultimately leads to the death of the insect [19,20]. This phase, during which the larvae are dying, also known as moribund, is characterised by exponential bacterial growth, a loss of larval body turgor and extensive melanisation [9,21].

Lifestyle switching — from a plant-beneficial root colonising to an insect-associated life — is an exciting new research line proposed not only for *P. protegens* but also for several other plant-beneficial rhizosphere bacteria such as *Streptomyces globisporus*, *Burkholderia* spp. and *Bacillus thuringiensis* [22]. Some of these bacteria were found to preserve their generalist, multi-host lifestyles whereas others are hypothesised to be the ancestors of highly specialised insect symbionts. There are several mechanisms that account for the evolutionary stability of such generalists. They are hypothesised to have an advantage over specialists in frequently changing environments [22–25]. Additional mechanisms help to maintain the sometimes large genetic repertoires required to survive in different environments. For example, a tight regulation of gene expression allows bacteria to reduce metabolic costs by only expressing factors when needed and the redundant use of factors preserves them across different environments [22,24]. The root environment is frequently changing, for example when plants grow and die, or when an insect feeds on plant material colonised by bacteria that are then instantly exposed to the insect gut environment. The type strain *P. protegens* CHA0, for instance, is one of the best studied bacteria experiencing such changes und exhibiting a multi-host lifestyle. A large regulatory network containing the Gac-Rsm pathway, allows CHA0 to sense changes in the environment and to adjust the expression of factors accordingly [26,27]. Indeed, Vesga et al. demonstrated that the transcriptome of CHA0 changed according to the colonised environment [15], and the insecticidal toxin Fit is an example of a compound which is produced in the insect’s haemolymph but not during plant root colonisation [20]. Notably, several factors are expressed in both root and insect environments, as demonstrated for some antimicrobial metabolites such as hydrogen cyanide, 2-4-diacetylphloroglucinol, and pyrrolnitrin. These compounds are presumably used to compete with microbial competitors in both environments [15]. Similarly, CHA0’s inositol catabolism genes are expressed during plant root and insect gut colonisation and demonstrated to contribute to root establishment [28].

Conservation of the multi-host lifestyle of *P. protegens* is corroborated by comparative analysis of closely related *P. protegens* strains obtained from diverse environments. The ability to colonise both plant and insect hosts, appears to be preserved across most *P. protegens* strains independent of their phylogenetic relationship and, to some extent, independent of their source of origin [12–14]. Isolates from different plants or insects were able to protect plants and kill insects [13], whereas only isolates from very distinct environments such as the human respiratory tract have lost their insecticidal activity at least partially [14]. Interestingly, all insect-derived *P. protegens* isolates described by Vesga et al. [13] exhibited strong insecticidal activity against *Plutella xylostella* larvae despite being isolated from healthy animals. Results obtained with CHA0 indicate that the ecological relationship with insects is even more complex. Depending on the insect species and the infection load, CHA0 either killed insect larvae or stayed associated with the insects until they reached adulthood and could eventually even use these adults as a vector for dispersal [21]. Furthermore, if fed to larvae of *Galleria mellonella*, CHA0 was rarely able to kill the larvae on its own but strongly benefited from a co-infection with entomopathogenic nematodes [29]. These examples highlight that, to date, many aspects of the ecology of the *P. protegens*-insect association remain unclear. The described findings suggest that CHA0, or the *P. protegens* species in general, are either differentially adapted to different insect species or are generally opportunistic rather than true insect pathogens [13,21,29].

The genetic characteristics of *P. protegens* and the conserved multi-host lifestyle observed throughout the species indicate that the multi-host lifestyle is not just a transition but an evolutionary stable state. Nevertheless, the evolutionary trajectories of bacterial-host interactions depend on many factors and are difficult to predict for single interactions [30]. Specialisation can be driven by environmental trade-offs but also when exposed to uniform environments [23–25]. The ability of bacteria to adapt to certain environments and the mechanisms underlying microbial adaptation can be uncovered by repeatedly propagating bacteria in defined environments and analysing resulting phenotypic and genotypic changes [31]. Using more complex environments such as *in vivo* passaging through a plant or an animal host can provide even more insights into how bacteria evolve in nature, e.g. how they adapt to new niches or hosts [32] as shown by various examples [33–37]. However, there are only few experiments where the adaptation of *Pseudomonas* spp. to multicellular organisms has been studied with respect to changes in bacterial virulence. Firstly, Jansen et al. [34] serially passaged the pathogen *Pseudomonas aeruginosa* PA14 through the nematode *Caenorhabditis elegans*. They found that adaptation was initiated through mutations in the Gac-Rsm global regulatory network which was followed by additional mutations leading to virulence attenuation. Loss of virulence was observed in most replicate lines independent of the immune status of the host [34]. Secondly, Meaden et al. [38] found that serially passaging *Pseudomonas syringae* pv. *tomato* PT23 on two different host plants led to an increase in colonisation ability of both plants only when evolved on the non-native host. However, the causal mutations for this change could not be identified [38]. Thirdly, Li et al. [39] serially passaged *P. protegens* CHA0 through a gnotobiotic plant root system. The loss of plant-detrimental effects in several lines was demonstrated to be connected to mutations in the Gac-Rsm network and subsequent loss of toxic secondary metabolite biosynthesis [39].

The major aim of this study was to investigate the robustness of the multi-host lifestyle of *P. protegens* CHA0 by testing whether the multifaceted properties of CHA0 would remain unchanged after a prolonged period of living as an insect pathogen. To this end, we recurrently exposed *P. protegens* CHA0 to non-sterile insect larvae and recorded strain- and population-level genotypes and phenotypes. Multiple independent CHA0 populations (hereafter also referred to as lines) were subjected to serial passage via repeated cycles of feeding to, and re-isolation from, larvae of the diamondback moth, *P. xylostella*. Selection of moribund larvae for re-isolation ensured that only CHA0 populations were transferred to the next cycle which were able to colonise and kill insect larvae. As a control for changes unrelated to the host, parallel bacterial populations were passaged in the absence of the insect host. We hypothesised that different evolutionary outcomes were conceivable. First, CHA0 could show little to no phenotypic or genotypic change if its genetic architecture constrains rapid niche specialisation which is suggested by the observed conservation of a multi-host lifestyle within the *P. protegens* species [12–14]. Second, CHA0 could evolve towards a more niche-specific lifestyle, as previously described for rhizosphere adaptation [39]. Such specialisation could manifest in increased insecticidal activity, enabling a more efficient exploitation of the host by accessing host-derived nutrients earlier. Additionally, specialisation towards an insect-associated lifestyle could lead to the loss of traits associated with its plant-beneficial, root-colonising lifestyle due to evolutionary trade-offs between these niches. Finally, by correlating phenotypic changes with genomic variation, we aimed to gain mechanistic insights into the regulatory networks and pathways that enable CHA0 to survive in and colonise insects.

## Results

### *Pseudomonas protegens* CHA0 remains insecticidal following serial passage

Ten independent replicate lines of *P. protegens* CHA0 were serially passaged through larvae of the insect pest *Plutella xylostella* (Fig 1). CHA0 was added to pellets of artificial food provided to the larvae in a non-sterile system. After three days of exposure, moribund larvae where surface-disinfected, homogenised, and plated onto selective medium to re-isolate CHA0. Colonies growing on the plates were pooled and this population was then used to inoculate the next batch of larvae. We refer to these lines as Ins4-13, where *Ins* stands for *Insect*. To evaluate potential adaptations of CHA0 to the background of the experimental system — specifically the feeding and re-isolation steps — three independent control lines were maintained in parallel. In these lines, referred to as Bkg1-3, where *Bkg* stands for *Background*, CHA0 was re-isolated directly from the food pellets without any contact with insect larvae.

**Fig 1:**
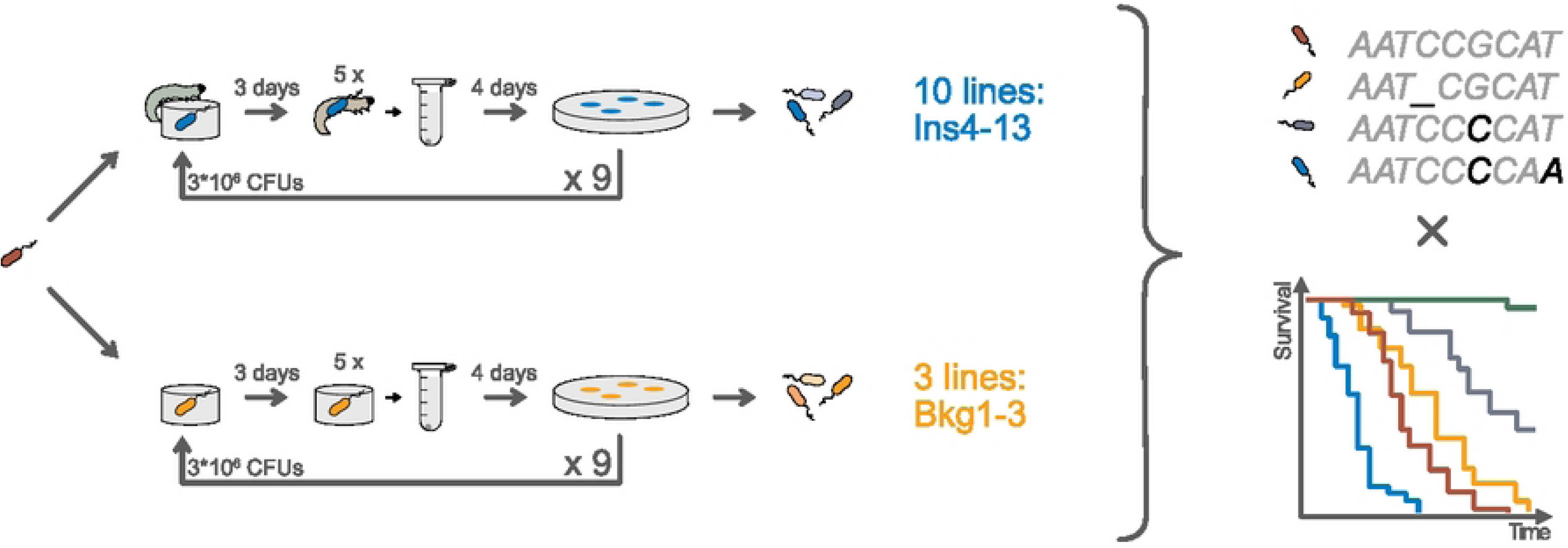
Overview serial passage experiment. Pseudomonas protegens CHA0 was serially passaged through larvae of the insect Plutella xylostella (independent replicate lines: Ins4-10) or through the background of the experimental system (independent replicate lines: Bkg1-3). For this, 3*10^6^ colony forming units (CFUs) of CHA0 were inoculated on pellets of artificial food and provided to second instar larvae in a non-sterile system. Five moribund larvae were collected after three days, surface-disinfected and homogenised. The homogenate was plated on selective agar to re-isolate CHA0 (see Fig S1.1A for detailed numbers of recovered bacteria). On average 4*10^4^ colonies were washed off the agar (see Fig S1.1B for detailed numbers of pooled colonies), and this population was used to infect the next batch of P. xylostella larvae. For the background lines, the same procedure was followed, but no larvae were added and CHA0 was re-isolated from five pooled food pellets. Each line was cycled ten times in total resulting in a minimum of 49 and 65 bacterial generations within the larvae or on the food pellet, respectively (see Fig S1.1C for detailed numbers of bacterial generations). All populations from cycles 1, 2, 4, 6, 8, and 10 as well as 12 single colonies picked per line from cycle 10 were sequenced and subjected to variant calling against the ancestor. Additionally, all populations from cycle 10 and some selected single colonies from cycle 10 were screened in different bioassays to compare to the ancestor.

After ten cycles, the re-isolated populations of each passaged line, were tested for changes in killing speed as an approximation of insecticidal activity and for colonisation of *P. xylostella* larvae (relative values see Fig 2, absolute values see Figs S2.1, S2.4A-B, individual experiments see Figs S2.2-3, S2.4C-D). No statistically significant changes could be observed when the insect and background passage treatments were compared to the original CHA0 (further referred to as ancestor) (Fig 2A, C). However, the three independent lines Bkg1, Ins5 and Ins8 killed the larvae slightly faster (Fig 2B) when individually compared to the ancestor. Moribund larvae were colonised on average with 6.4*10^5^ colony forming units (CFUs)/larva by the ancestor (Fig S2.4A). Relative colonisation did not change, neither for the passage treatments (Fig 2E) nor for the individual independent lines (Fig 2F). However, lines Ins5 and Ins8 showed a tendency towards increased colonisation levels, indicating that a faster colonisation might have led to the observed faster killing speed in these lines. This pattern could not be observed for line Bkg1.

**Fig 2:**
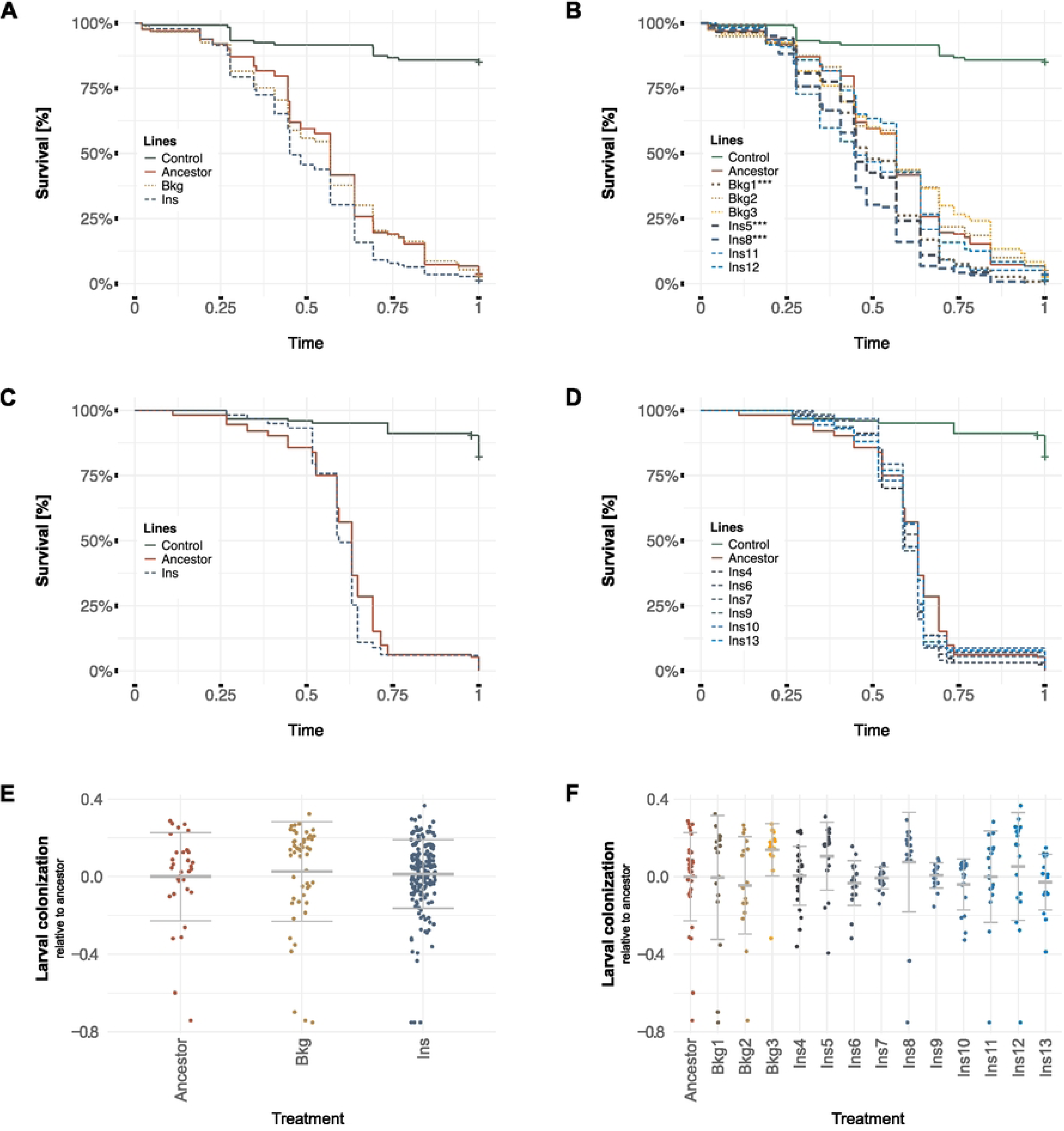
Ability of serially passaged Pseudomonas protegens CHA0 to kill and colonise larvae of Plutella xylostella. **A-D** Survival of P. xylostella larvae infected with the different insect and background lines over relative time, until 95% of larvae were dead, is depicted as Kaplan-Meier curves. Lines were tested in two batches: **A**, **B** batch 1 and **C**, **D** batch 2. Absolute times can be found in Fig S2.1. **E**, **F** Relative colonisation of moribund larvae by the different lines. Plots show individual data points, mean and standard deviation. Colony forming units (CFUs) were determined by plating individual, homogenised larvae on selective medium. Relative values were calculated by dividing the treatment value by the mean of the values obtained for the ancestor in the respective experimental repetition. Absolute values can be found in Fig S2.4A-B. **A**, **C**, **E** Results for the insect passage treatment (Ins) and the background passage treatment (Bkg). **B**, **D**, **F** Results for individual independent lines (Bkg1-3, Ins4-13). Survival and colonisation data represent pooled data from three independent experiments (N=3) per line with 42 (survival) and six (colonisation) larvae per individual line and repetition. Statistical analysis was performed with the pooled data sets. Lines Bkg1, Ins5, Ins8 significantly differed from the ancestral CHA0 in killing speed and are indicated with asterisks (*** = P < 0.001). Individual experiments are shown in Figs S2.2-3, S2.4C-D.

Overall, insecticidal activity and larval colonisation did not change substantially after passaging CHA0 for ten cycles, neither when having the opportunity to adapt to insect larvae nor when passaged through the background only, although three independent lines exhibited a slight increase in killing speed.

### Population-level mutations are associated with nutrient acquisition and energy metabolism

After the 1^st^, 2^nd^, 4^th^, 6^th^, 8^th^, and 10^th^ cycle, the re-isolated populations were investigated for genomic changes relative to the ancestor using deep whole-genome shotgun sequencing, yielding an average genome-wide coverage of approximately 340 per population. For all samples, more than 99% of the reads mapped to the CHA0 reference genome, and the absence of consistently annotated sequences among unmapped reads confirmed that no contaminants were present in the selection procedure.

Variant calling identified a total of 33 mutations across all cycles sequenced including single nucleotide polymorphisms, small insertions and deletions, of which 32 occurred at unique base pair positions. A complete list of mutations, along with their corresponding locus tags and putative functions, is provided in Table 1. The majority of mutations (29 out of 33) were located within coding regions. Of these, two coding regions were hit in more than one line: *PPRCHA0_1039* annotated as a LysR type transcriptional regulator (LTTR) and *PPRCHA0_2684* annotated as a putative sensor protein RstB. *PPRCHA0_1039* was mutated in three independent lines, each carrying a different mutation (*PPRCHA0_1039*^Leu21fs, Arg32fs, Gln260*^). This region was hit in insect (Ins12) as well as background lines (Bkg2 and Bkg3) indicating that the mutations were driven by the background of the experimental system rather than by the interaction with the insect larvae. Notably, the mutation *PPRCHA0_2684*^Gly389Cys^ occurred at the exact same base pair in two different independent lines (Bkg3 and Ins7). All coding regions hit by a mutation were subjected to gene set enrichment analysis (Fig 3A). This analysis revealed a significant overrepresentation of COG (Clusters of Orthologous Genes) category G (Carbohydrate transport and metabolism) followed by category E (Amino acid transport and metabolism). These findings indicate that nutrient acquisition and energy metabolism were primary targets of selection within the passage system. No specific KEGG (Kyoto Encyclopedia of Genes and Genomes) pathways were enriched (Fig S3.1).

**Fig 3:**
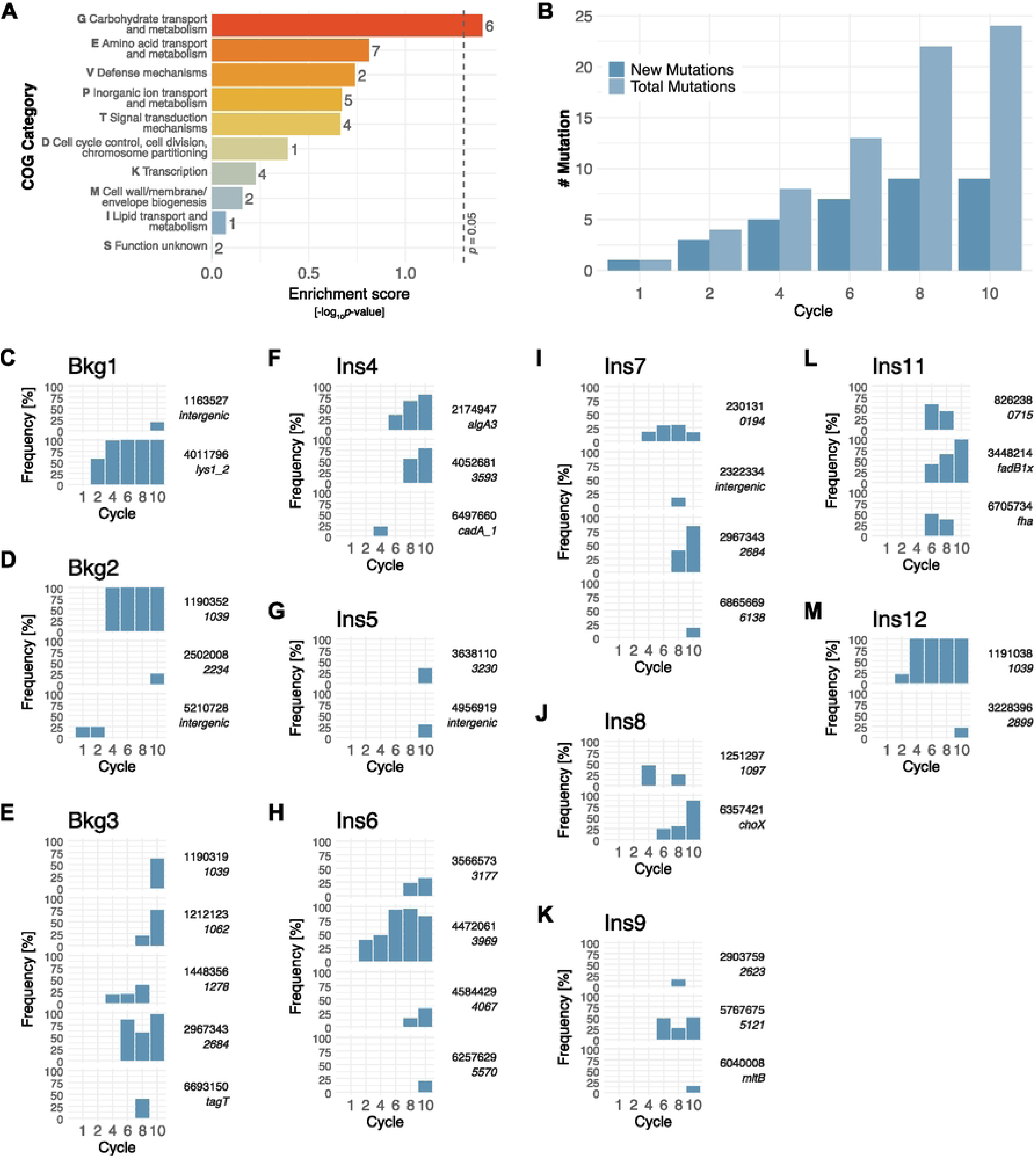
Population level mutations found in the serially passaged lines. **A** COG (Clusters of Orthologous Genes) categories enrichment analysis of all coding regions hit by a mutation. For KEGG (Kyoto Encyclopedia of Genes and Genomes) pathway enrichment see Fig S3.1. **B** Cumulative numbers of newly arising and total mutations (single nucleotide polymorphisms and small insertions and deletions) per cycle. **C**-**M** Frequencies of individual mutations over cycles are shown per background (Bkg1-3) or insect (Ins4-13) line. Exact base pair positions as well as gene names or if not available locus tags are displayed.

**Table 1:**
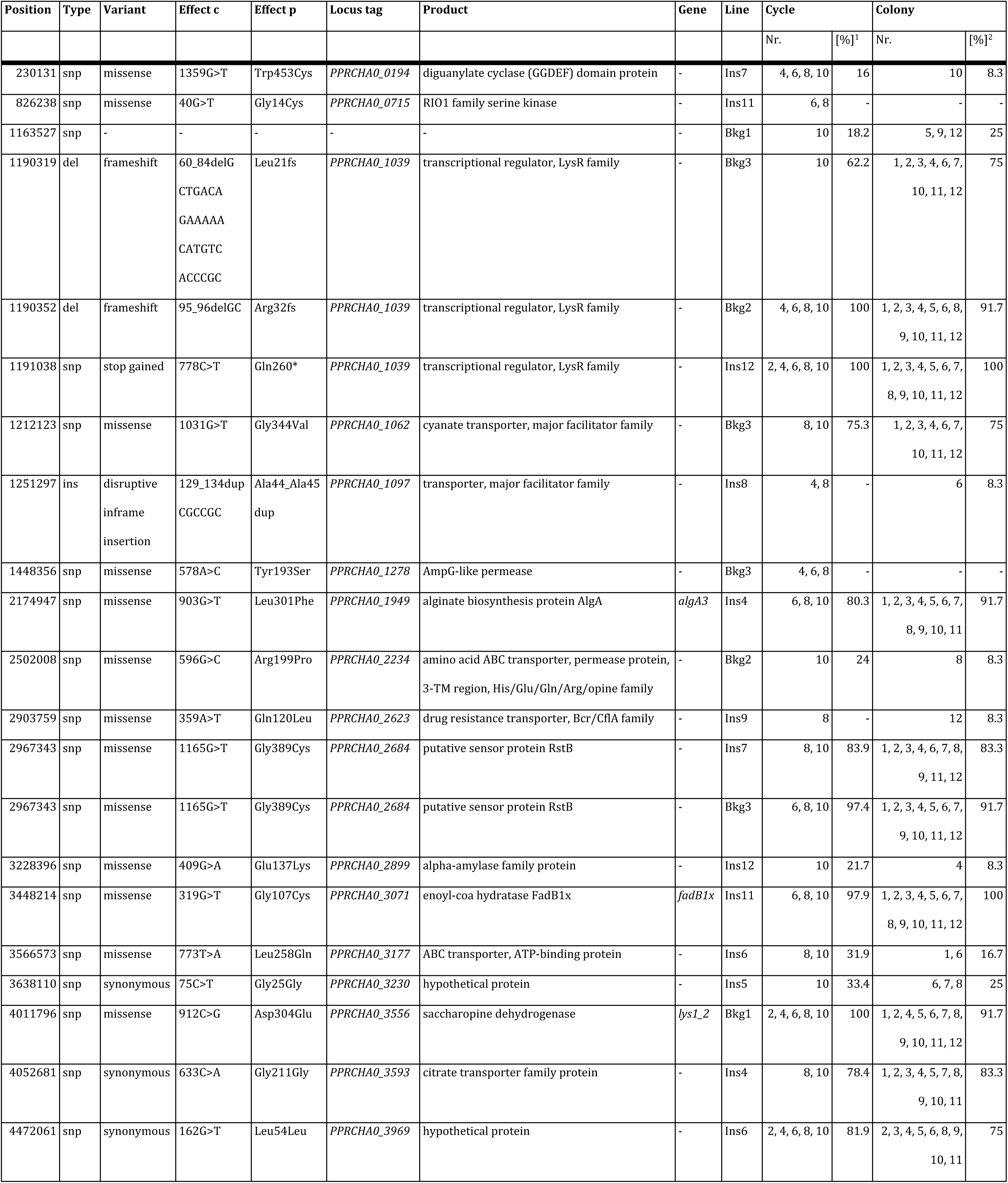
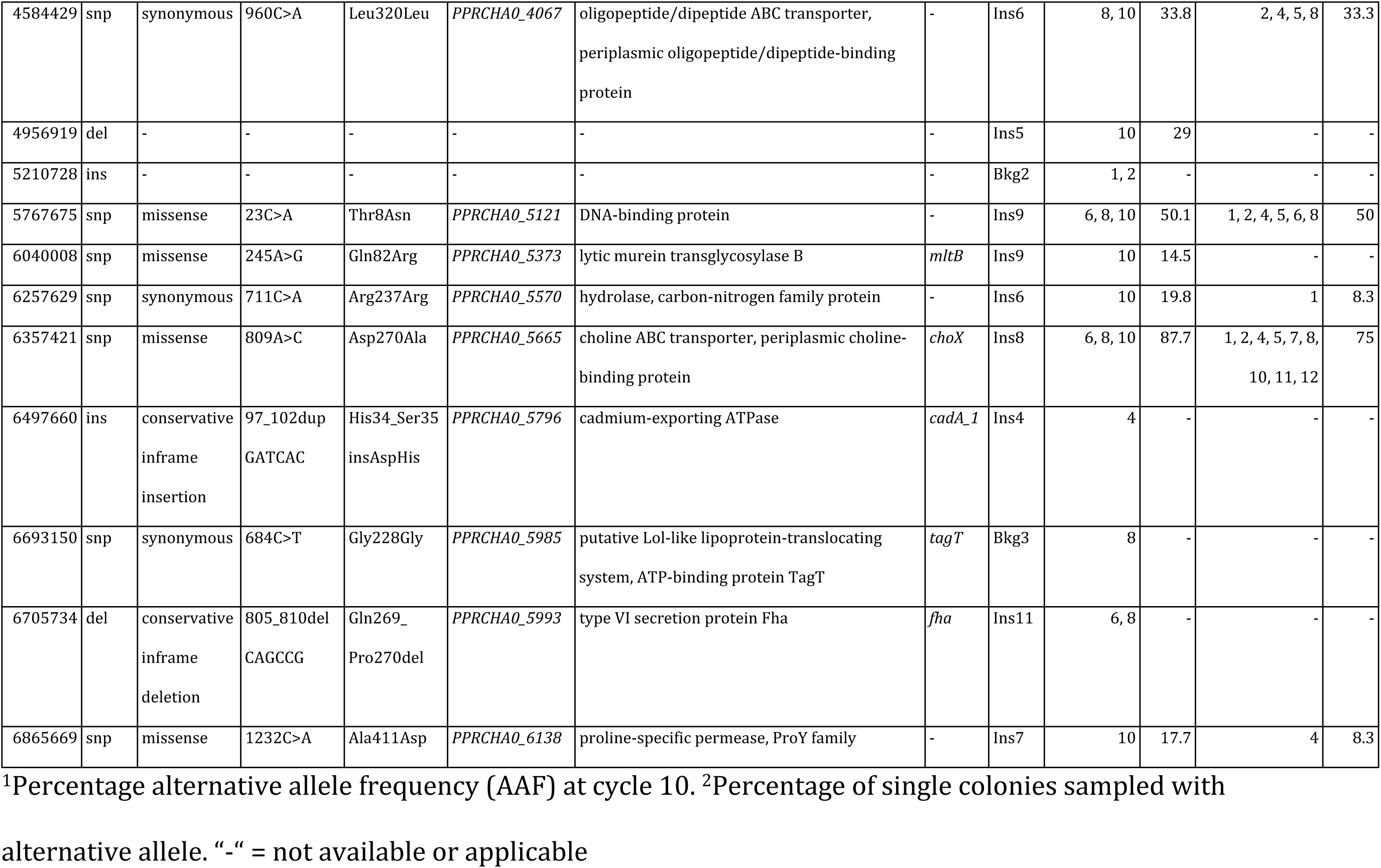
Mutations found in background and insect lines.

The number of total and newly appearing mutations per cycle increased until the last cycle, indicating that an equilibrium was not yet reached (Fig 3B). Some of the mutations increased in frequency and persisted until cycle 10 whereas others disappeared again (Fig 3C-M). 19 mutations were detected in at least two consecutive time points and mutations with a frequency of *>* 50% in cycle 10 included *PPRCHA0_1062*^Gly344Val^, *algA3*^Leu301Phe^, *fadB1x*^Gly107Cys^, *lys1_2*^Asp304Glu^*, PPRCHA0_5121*^Thr8Asn^, *choX*^Asp270Ala^, and the synonymous mutations in *PPRCHA_03593*^Gly211Gly^ and *PPRCHA0_3969*^Leu54Leu^ (Fig 3C-M, Table 1). In general, several different permeases and transporters (*PPRCHA0_1062*, *PPRCHA0_1097*, *PPRCHA0_1278*, *PPRCHA0_2234*, *PPRCHA0_2623*, *PPRCHA0_3177*, *PPRCHA0_3593*, *PPRCHA0_4067*, *choX*, *cadA_1*, *PPRCHA0_6138*) were hit by mutations, some of them becoming fixed, whereas others disappeared again (Table 1). These coding regions were mostly associated with the COG categories E and G, and probably resulted in the above described enrichment (Fig 3A).

In addition to the populations, 12 single colonies were picked per line after cycle 10 and investigated for strain-level genomic changes. These strains carried most of the mutations found in the respective populations, although some additional mutations were detected (see Table ST1.1 for all mutations found).

Taken together, most mutations occurred at different coding regions although an enrichment for carbohydrate transport and metabolism was reported. No parallelism for one genomic region was observed for insect adaptation, whereas at least one region seemed to be under selection by the background of the system.

### Few lines changed in antimicrobial, swarming and growth characteristics

Serially passaged insect and background lines were further phenotypically characterised at the population-level after the tenth cycle for several plant- and insect-associated traits in addition to insecticidal activity and larval colonisation. To colonise plant roots, *P. protegens* CHA0 requires different rhizosphere competence traits such as motility, biofilm formation and antimicrobial competition [40]. The antimicrobial abilities of CHA0 also result in the inhibition of different plant pathogenic oomycetes and fungi [4,8]. Consequently, swarming motility, biofilm formation and direct fungal and oomycete inhibition were tested in different *in vitro* setups whereas plant root colonisation and protection were tested *in planta*. Furthermore, we assessed the ability of CHA0 to compete with bacteria it might encounter while infecting insects [16], its behaviour when exposed to stressors found in the insect gut such as pH differences and oxidative stress [41], and its ability to grow in haemolymph mimicking media in different *in vitro* setups. For all bioassays, relative values were calculated by dividing the treatment value by the mean of the values obtained for the corresponding ancestral CHA0 strain in the respective experiment (Fig 4Fig 4; for absolute values, individual experiment and additional relative values see Figs S4.1-19). No statistically significant differences were observed in any bioassay when comparing the insect or background passage treatments to the ancestor or to each other. Only the biofilm formation ability was higher in the background than in the insect passage treatment. Altogether, this indicates that none of the serial passage regimes had a major effect on the phenotypes of CHA0. We did, however, detect some subtle changes in individual independent replicate lines in comparison to the ancestor, as described in the following paragraphs.

**Fig 4:**
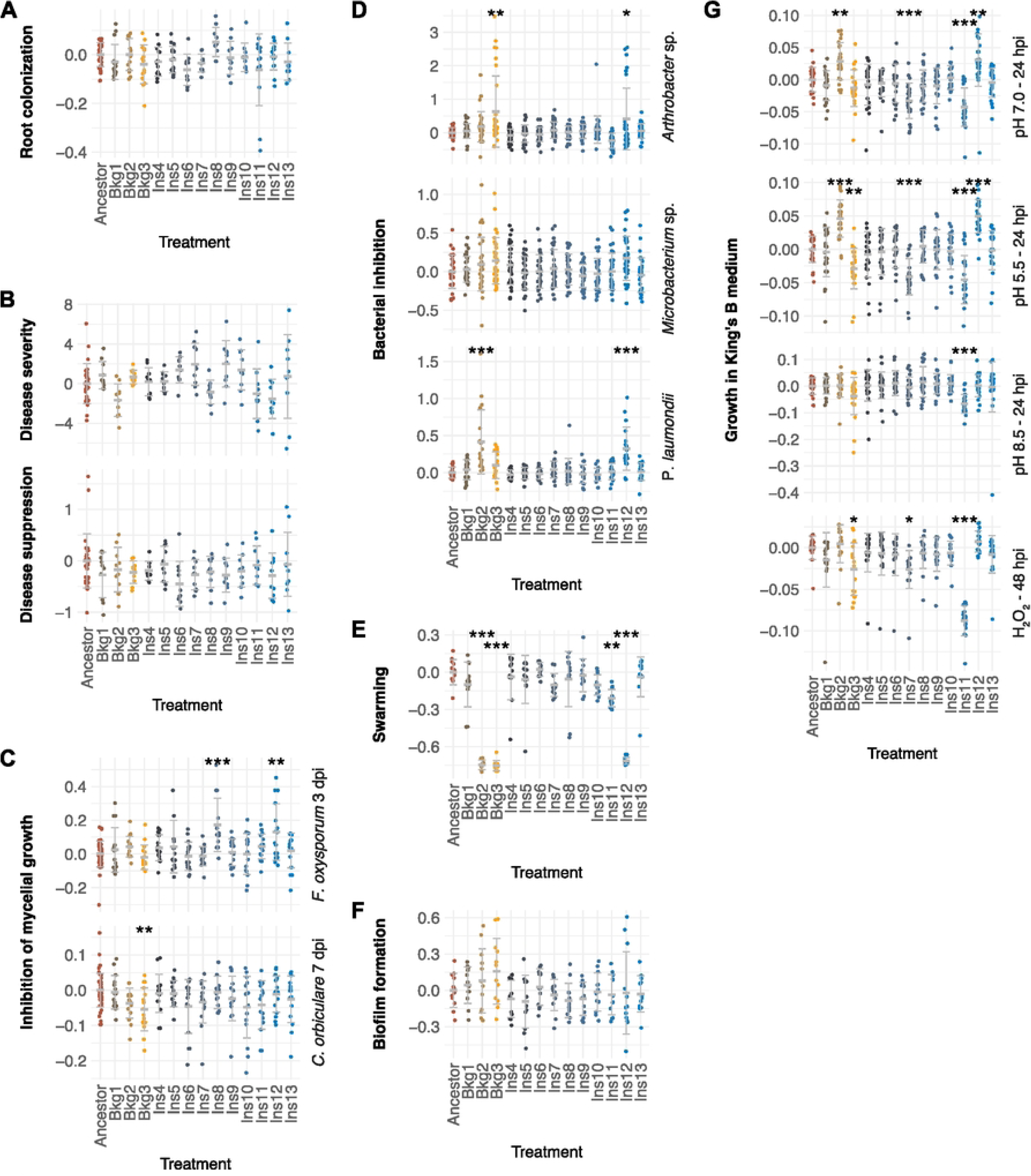
Phenotypic characteristics of serially passaged CHA0 in comparison to the ancestral CHA0. All plots show relative values of the individual independent lines in comparison to the ancestor. **A, B** Cucumber seedlings were planted in non-sterile soil with or without the plant pathogen Pythium ultimum Pu-11 and drench inoculated with the respective bacterial treatments. Plants were harvested at 11 days post infection (dpi). **A** Root colonisation of healthy cucumber plants by the different lines was assessed as colony forming units (CFUs) per g root. **B** Disease severity was calculated as percentage shoot weight loss with the pathogen divided by the respective bacterial treatment without the pathogen. Disease suppression was calculated as percentage disease severity in the bacterial treatment compared to the disease severity in the uninoculated control. **C** Inhibition of the fungal plant pathogens Fusarium oxysporum f. sp. radicis-lycopersici FORL22 and Colletotrichum orbiculare D323 was assessed in vitro on malt agar and determined as percentage of mycelial diameter reduction in the presence of bacteria compared to controls without bacterial treatments. **D** Inhibition of the Plutella xylostella gut isolates Arthrobacter sp. Px40 and Microbacterium sp. Px99, and the insect pathogen Photorhabdus laumondii DJC was assessed in vitro on lysogeny broth (LB) agar by measuring inhibition zone size at 2 dpi. **E** Swarming ability was assessed on 0.4% agar swarming plates by measuring the swarm diameter at 20 hours post inoculation (hpi). **F** Biofilm formation in one-third strength King’s B medium(1/3 strength KB) using a 96-well plate system with peg lids was assessed after 18 h indirectly by staining the biofilm with crystal violet and subsequent optical density measurement of the washed off stain at 590 nm (OD_590_). **G** Growth of the lines in 1/3 strength KB (KB pH 7.0) was assessed by measuring optical density at 600 nm (OD_600_) at different time points. In 1/3 strength KB, bacteria were additionally exposed to pH (KB pH 5.5, KB pH 8.5) or oxidative stress (0.001% H_2_O_2_ (KB H_2_O_2_)). All plots depict individual data points, mean and standard deviation based on pooled data from two to five independent experiments (N=2-5). Statistical analysis was performed with the pooled relative data and asterisks indicate statistical differences to the ancestor (*, **, *** = P < 0.05, 0.01, 0.001). Additional bioassay data, absolute values, individual experiments and data for the insect and background passage treatments are shown in Figs S4.1-19.

Plant root colonisation and plant protection efficiency were assessed in a *Pythium ultimum*-cucumber system. For this purpose, cucumber seedlings were planted in non-sterile soil with or without the plant pathogenic oomycete *P. ultimum* Pu-11 (see Table 2) and drench-inoculated with the respective bacterial treatments. Plants were harvested at 11 days post infection (dpi) to determine bacterial root colonisation and the impact of the bacterial treatments on plant growth, disease severity and disease suppression (Fig 4A-B, S4.1-5). The ancestor colonised healthy roots at an average density of 4.2*10^6^ CFUs/g root and increased shoot fresh weight by 8.1%. The independent lines showed no differences in relative root colonisation or plant growth in comparison to the ancestor (Fig 4A), although line Ins11 displayed lower colonisation in some outliers. Uninoculated control plants exhibited a mean disease severity of 85.8%, while those inoculated with the ancestor only reached 10.6% disease severity, corresponding to 91.9% disease suppression. Plant protection efficiency was preserved in all lines, except for Ins6, which showed a tendency towards increased relative disease severity and reduced relative disease suppression compared to the ancestor (Fig 4B).

**Table 2:**
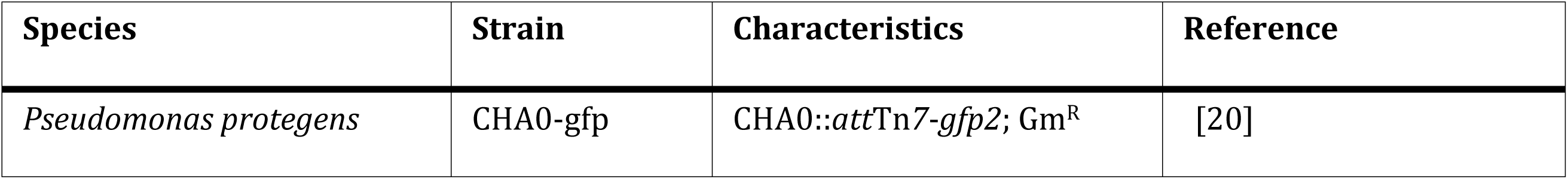
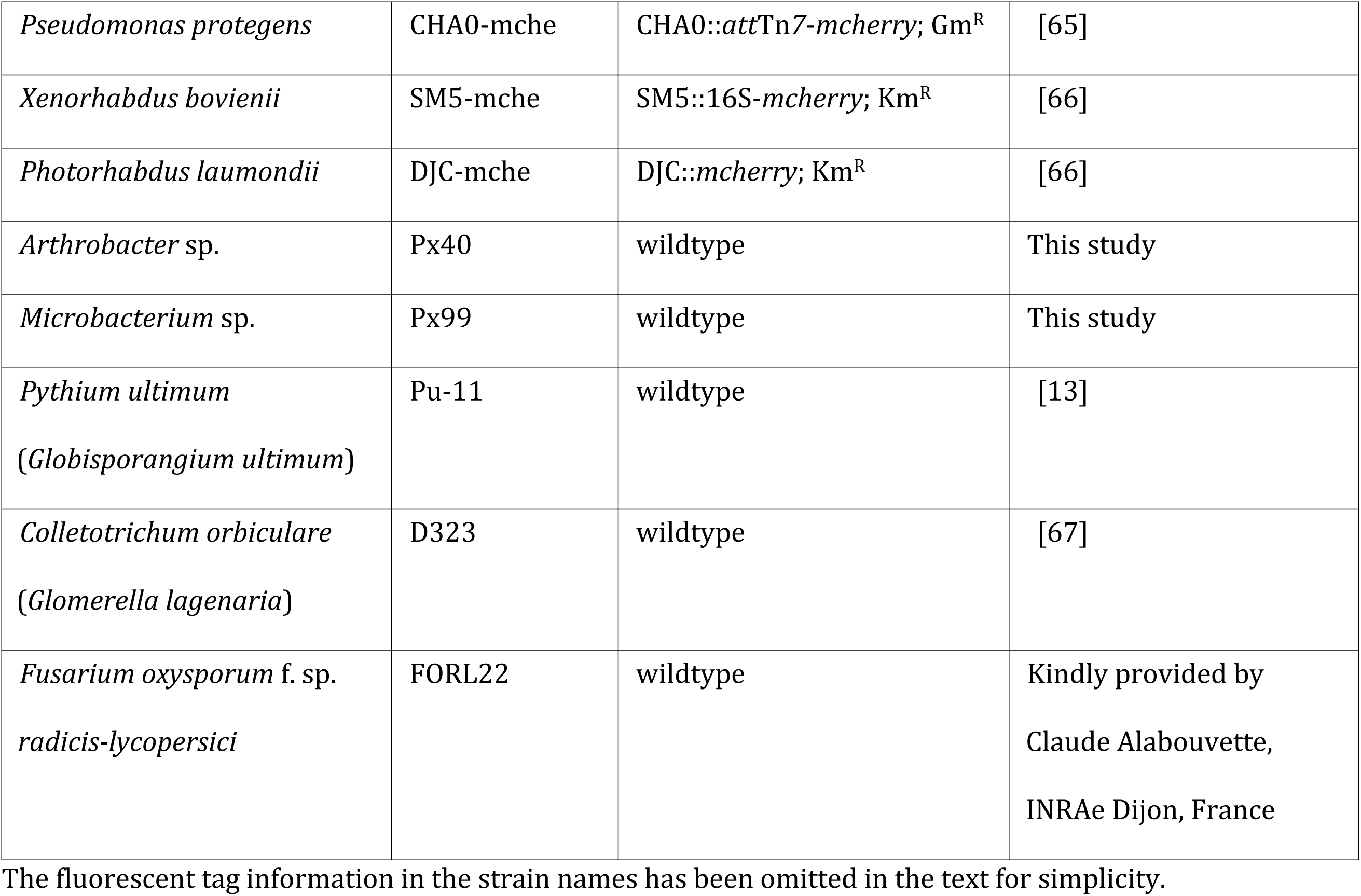
Microorganisms used in this study.

In addition to direct and indirect effects on plants, we investigated the ability of the lines to inhibit the growth of the following microorganisms on agar plates: the plant pathogenic fungi *Fusarium oxysporum* f. sp. *radicis-lycopersici* FORL22 and *Colletotrichum orbiculare* D323, the plant pathogenic oomycete *P. ultimum* Pu-11, the insect gut isolates *Arthrobacter* sp. Px40 and *Microbacterium* sp. Px99 as well as the insect pathogenic bacteria *Photorhabdus laumondii* DJC and *Xenorhabdus bovienii* SM5 (see Table 2). Fungal inhibition was estimated as radial mycelial growth reduction whereas inhibition zone size was compared for the bacterial competitions. Mean percentage inhibition of *F. oxysporum* by the ancestor increased from 34.6% at 3 days post inoculation (dpi) to 60.5% at 7 dpi (Fig S4.6). Lines Ins8 and Ins12 showed an increase in relative inhibition of *F. oxysporum* at 3 dpi meaning that these lines were better in restricting the fungal growth than the ancestor (Fig 4C). The effect was preserved from 3 to 7 dpi for Ins8 whereas Ins12 only exhibited a similar tendency at the later time point (Fig S4.9). Mean percentage inhibition of *C. orbiculare* by the ancestor remained at similar levels from 48.5% at 7 dpi to 49.1% at 14 dpi (Fig S4.6). Bkg3 showed a decrease in relative inhibition at 7 dpi, but the effect was not preserved until 14 dpi (Fig 4C, S4.9). At 14 dpi, Ins12 displayed a tendency towards increased relative inhibition (Fig S4.9). Mean percentage inhibition of *P. ultimum* by the ancestor was 31.7% at 3 dpi and relative inhibition did not change for any of the independent lines (Fig S4.9) which is in line with the findings in the *P. ultimum*-cucumber assays. At 2 dpi, the ancestor caused mean inhibition zone sizes of 55.8 px for *Arthrobacter* sp. Px40, 31.0 px for *Microbacterium* sp. Px99, 59.4 px for *P. laumondii*, and 55.7 px for *X. bovienii* (Fig S4.10), with 1 mm corresponding roughly to 11.5 px. Bkg2, Bkg3 and Ins12 displayed an increase in relative inhibition of the *Arthrobacter* strain and/or *P. laumondii* (Fig 4D). No significant changes were observed with the other two bacteria *Microbacterium* sp. 99 and *X. bovienii* (Fig 4D, S4.13).

Finally, we investigated different *in vitro* phenotypic characteristics including swarming ability, biofilm formation, sensitivity to pH and oxidative stress. The swarm diameter measured on soft agar was 30.3 mm for the ancestor (Fig S4.14) and a pronounced decrease in relative swarming ability was found for Bkg2, Bkg3, Ins11, and Ins12 (Fig 4E). Biofilm formation in one-third strength King’s B medium (1/3 strength KB) using a 96-well plate system with peg lids was measured after 18 h. Absorbed crystal violet resulted in a mean optical density at 590 nm (OD_590_) of 1.80 for the ancestor (Fig S4.15). None of the lines showed any differences in relative biofilm formation, indicating that biofilm formation was not affected by the serial passage regimes (Fig 4F). Growth was measured in the haemolymph-mimicking Grace’s insect medium (GIM), in 1/3 strength KB at neutral pH (KB pH 7.0, same as in the passage experiment), when challenged with oxidative stress (KB 0.001% H_2_O_2_), and at high and low pH stress (KB pH 4.5 and KB pH 8.5). Optical density at 600 nm (OD_600_) was used as an approximation for growth, however, it should be noted that this method does not distinguish between viable cells and non-viable cells or extracellular matrix. OD_600_ was measured at three time points: at late exponential phase around 14-17 hours post inoculation (hpi), at early stationary phase around 24 hpi and at late stationary phase around 48 hpi (Fig 4G, S4.19). At 48 hpi, the ancestor grew to an OD_600_ of 0.98 in GIM and 1.71 in KB pH 7.0 displaying a similar growth with the diverse stresses tested. In GIM, no changes were reported for any of the passaged lines. Line Ins11 showed a clear growth deficiency in KB pH 7.0 and in all stresses studied at 24 and 48 hpi. Bkg2, Bkg3, Ins7, and Ins12, were differentially affected in their relative growth in KB pH 7.0 and under pH and oxidative stress. The decreased growth of lines Ins5, Ins8, and Ins10 in KB H_2_O_2_ was observed in only one out of three replicates.

Some selected single colonies were characterised for strain-level changes in bacterial inhibition and growth characteristics (Figs S4.20-25). The results were mostly comparable to the populations if the strains carried the same mutations.

Taken together, the ability to colonise plant roots, protect the plants from *P. ultimum*, and form biofilms was preserved in all individual independent background and insect lines. Phenotypic changes regarding *in vitro* antimicrobial activity, swarming ability, and growth occurred in some insect as well as in background lines.

### Mutations in specific regulatory systems and fatty acid metabolism underlie phenotypic shifts

To investigate potential mechanisms underlying the observed phenotypic changes detected in some of the lines, we compared the occurrence of mutations and changes in the bioassays (Fig 5, S5.1A-B) and tested for statistically significant correlations (Fig S5.1C). Lines Bkg2, Bkg3, and Ins12 each had a mutation in *PPRCHA0_1039* encoding a LTTR with alternative allele frequencies (AAF) at cycle 10 of 100%, 62.2% and 100%, respectively. LTTRs consist of a DNA recognition helix and a regulatory domain containing a signal binding cavity [42]. The Arg32fs and Leu21fs mutations observed in Bkg2 and Bkg3, respectively, resulted in frameshifts towards the middle or end of the DNA recognition helix. This led in both cases to the introduction of early stop codons and consequently shortened, likely non-functional peptides (Fig S7.1). The Gln260* mutation identified in Ins12 introduced a stop codon in the C-terminal regulatory domain of the protein, potentially leading to an altered tetrameric 3D protein structure as evidenced by structural analysis using AlphaFold (Fig S7.1). Consequently, signal recognition via the LTTR *PPRCHA0_1039* is likely disrupted in all three cases. Bkg2 and Ins12 exhibited an increase in antimicrobial activity against *Arthrobacter* sp. 40 and *P. laumondii*, a decrease in swarming ability and an increase in growth in 1/3 strength KB with and without certain stressors. Bkg3 only displayed part of these characteristics but additionally showed a decrease in antimicrobial activity against *C. orbiculare*. The frequency of the mutations in *PPRCHA0_1039* in the different populations significantly correlated with relative swarming ability and antimicrobial activity against *P. laumondii* (Fig S5.1C).

**Fig 5:**
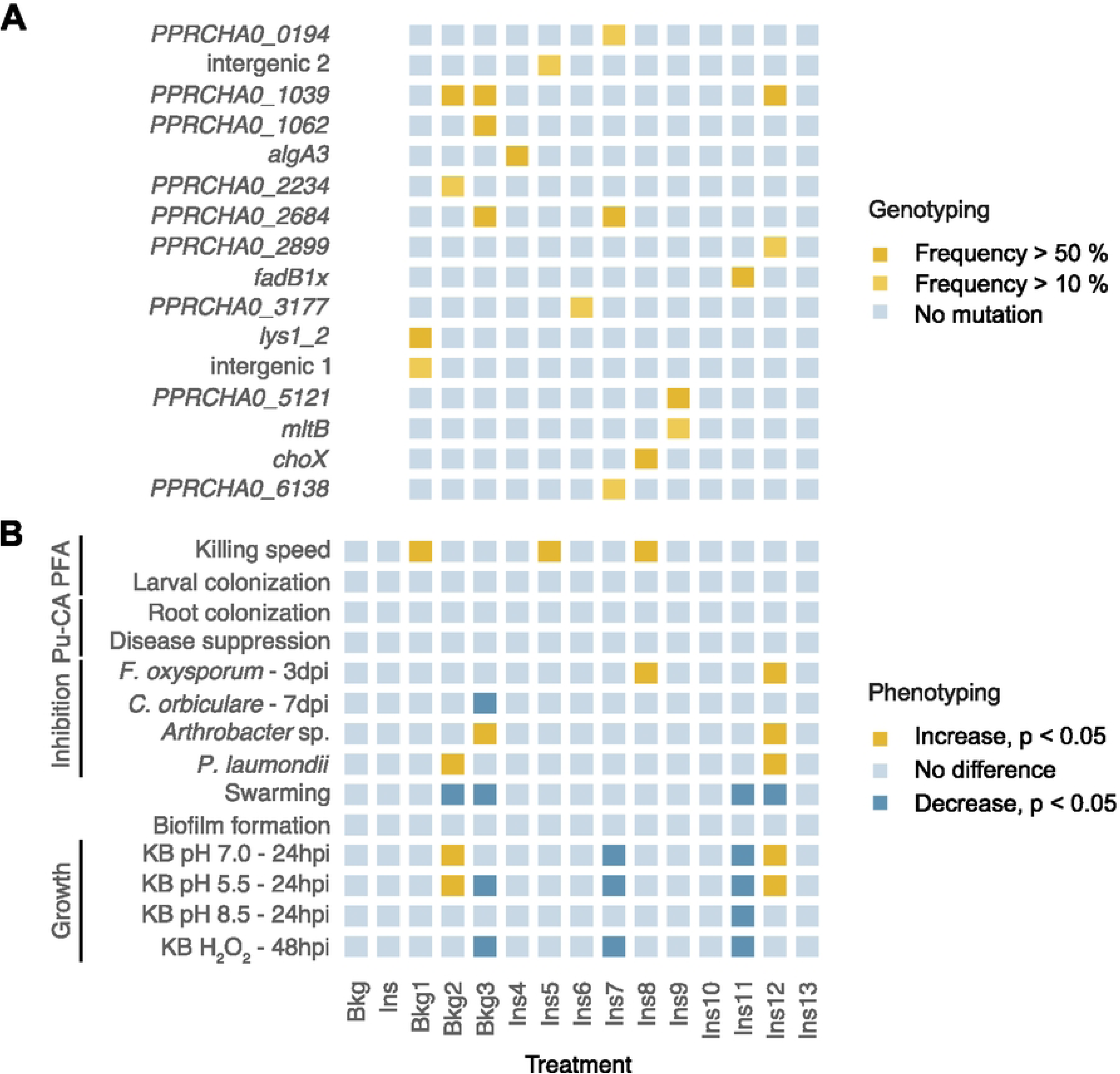
Overview on phenotypes and genotypes of different lines. The occurrence of mutations was compared with changes in the bioassays. **A** Genomic regions (gene names or locus tags) of each line with a mutation present at cycle 10 with more than 10% frequency. **B** Selected bioassays resulting in significant in- or decreases of passage treatment (Bkg, Ins) or individual independent lines (Bkg1-3, Ins4-13) in comparison to the ancestor. For all bioassays and the complete correlation matrix see Fig S5.1. Abbreviations: PFA = Plutella xylostella feeding assay, Pu-CA = Pythium ultimum-cucumber assay, KB = one-third strength King’s B medium at pH 7, 5.5 or 8.5 or supplemented with 0.001% H_2_O_2_, dpi = days post inoculation, hpi = hours post inoculation.

The putative sensor protein RstB encoding region *PPRCHA0_2684* was hit with the same mutation in Bkg3 and Ins7 with an AAF at cycle 10 of 97.4% and 83.9%, respectively. This missense mutation — potentially changing the interaction of the protein with ATP in the catalytic domain (Fig S7.2) — might interfere with downstream signal transduction and could contribute to fitness in the serial passage experiment [43]. Ins7, containing the *PPRCHA0_2684*, but not the *PPRCHA0_1039* mutation, displayed a reduction in growth in 1/3 strength KB with and without stressors, while Bkg3, having both mutations, only showed part of these characteristics.

Ins11 was growth deficient in 1/3 strength KB with and without stressors. This line has a mutation, with an AAF at cycle 10 of 97.9%, in the region encoding the substrate binding site of fadB1x. *fadB1x* encodes an enoyl-coa hydratase involved in fatty acid metabolism. The mutation likely impacts the protein’s function (for predicted changes in 3D structure see Fig S7.3), thus resulting in the observed growth deficiency and corroborated by the statistically significant correlations between the mutation and *in vitro* growth characteristics (Fig S5.1C). Additionally, Ins11 showed a tendency towards decreased root colonisation and decreased antimicrobial activity against *C. orbiculare* and *Arthrobacter* sp. Px40 which might be a result of the reduced growth.

All lines with an increase in killing speed had at least one mutation in a coding region. Those were in *lys1_2* (AAF at cycle 10: 100%) encoding a saccharopine dehydrogenase for Bkg1, the hypothetical gene *PPRCHA0_3230* for Ins5 (AAF at cycle 10: 33.4%) and *choX* (AAF at cycle 10: 87.7%) encoding the periplasmic binding protein of a choline transmembrane transporter for Ins8. The mutation in *lys1_2* led to a replacement of asparagin with glutamine which did not affect the 3D structure (Fig S7.4), whereas the mutation in the putative gene *PPRCHA0_3230* was synonymous. The mutation in *choX* resulted in the exchange of asparagine with alanine towards the end of the protein. This potentially changes the surface of the protein in proximity to the substrate binding site (Fig S7.5). For each, Bkg1 and Ins5, another non-coding region was additionally hit by a mutation with low alternative allele frequencies at cycle 10. For Bkg1, the mutation was between a hypothetical gene and *algA1* and for Ins5 between *rffA* and a putative lipoprotein.

In summary, some phenotypes could be associated with certain mutations, especially within *PPRCHA0_1039* and *fadB1x*. However, it is important to note that several of the above discussed lines have additional mutations, which could impact their phenotypes, too.

### *Pseudomonas protegens* CHA0 does not undergo strong bottlenecks during insect infection

Bottlenecks affecting CHA0 during survival on food pellets and larval infection might affect the outcome of the serial passage experiment. We therefore investigated the severity of a potential bottleneck in our system using a mixture of differentially fluorophore-tagged *P. protegens* CHA0 variants. Specifically, mCherry-expressing CHA0-mche was mixed with GFP-expressing CHA0-gfp in defined initial ratios: 1:1, 1:10^1^, 1:10^2^, 1:10^3^,1:10^4^. These mixtures were added to pellets of artificial food and provided to *P. xylostella* larvae in the same system as used before. This resulted in 51±5 CHA0-gfp cells added to the pellets for the 1:10^4^ treatment.

The number of CHA0-mche and CHA0-gfp CFUs was determined for all treatments, and in most cases, both strains could be recovered (Fig 6A). Only the 1’000 or 10’000-fold dilutions of CHA0-gfp in the 1:10^3^ and 1:10^4^ treatments led to its loss, meaning no CHA0-gfp was detected above the detection limit, in 2/16 and 2/8 larvae, respectively. CHA0-gfp could be recovered in all cases from the pellets, even in the 1:10^4^ treatment.

**Fig 6:**
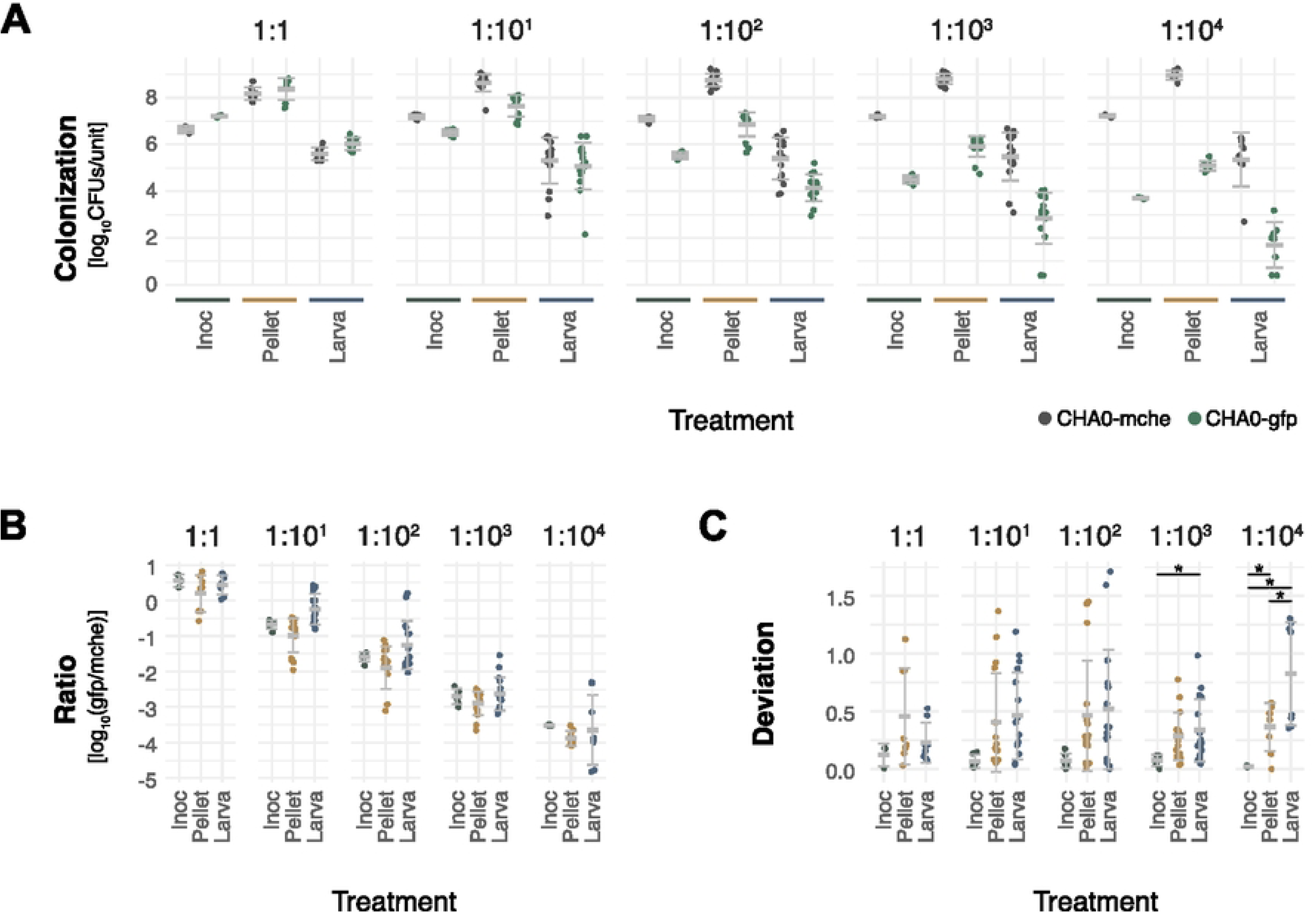
Pseudomonas protegens CHA0 tagged with different fluorophores was used to determine bottleneck severity. CHA0 expressing mCherry (CHA0-mche) was mixed with CHA0 expressing GFP (CHA0-gfp) in different initial ratios: 1:1, 1:10^1^, 1:10^2^, 1:10^3^, 1:10^4^. 1.7*10^5^ colony forming units (CFUs) were used on average to inoculate pellets of artificial food without or with larvae of Plutella xylostella to feed on it. CFUs of CHA0-mche and CHA0-gfp were determined in the inoculums, and at 30 hours post infection (hpi) in pellets and surface-disinfected larvae. **A** Recovered absolute numbers of CFUs for inoculum [CFUs/ml], pellet [CFUs/pellet] or larva [CFUs/larva], **B** log_10_(CHA0-gfp/CHA0-mche), and **C** the deviation from the mean ratio in the inoculum are displayed for the different treatments. Single replicates, mean and standard deviation of pooled data from two independent experiments (N=2 for Neg, 1:10^1^, 1:10^2^, 1:10^3^ and N=1 for 1:1, 1:10^4^, six replicates per treatment and experiment) are shown. Statistical analysis was performed with the pooled deviation data and asterisks indicate statistical differences (*, **, *** = P < 0.05, 0.01, 0.001).

The recovered ratios in the inoculums were on average comparable with the ratios from the pellets and the larvae (Fig 6B), indicating that none of the strains had an unwanted fitness advantage or disadvantage. A deviance from the expected inoculum ratio (Fig 6C) indicates that the populations in these samples did undergo a bottleneck. For larvae, the 1:10^3^ and 1:10^4^ treatments were affected whereas for the pellets only the 1:10^4^ treatment was statistically significantly deviating. Together, these observations indicate that bottlenecks did not play a major role in the serial passage experiment. The effect was higher for larvae infection then for growth on the food pellets.

## Discussion

### *Pseudomonas protegens* CHA0 preserves its insect- and plant-associated lifestyles

In this study, we investigated whether serial passaging of *P. protegens* CHA0 through larvae of the diamondback moth *Plutella xylostella* via repeated cycles of colonisation and killing of said larvae would result in a more insect-adapted lifestyle of the bacteria. To control for adaptations caused by the serial passage system itself, background lines were passaged in the same system but in absence of the insect host. We proposed two competing hypotheses. First, there was the possibility that CHA0 preserves its multi-host lifestyle and does not adapt rapidly. Second, CHA0 could adapt a more insect-associated lifestyle which could result in the loss of plant-associated traits. Adaptation towards the insect larvae was hypothesised to lead to increased virulence of CHA0, as moribund larvae were selected for during the serial passage. To evaluate robustness of CHA0’s multi-host lifestyle, we tested among other things virulence towards insects by measuring both the level of colonisation and the speed of killing, evaluated plant interactions by root colonisation quantification and plant protection efficiency, and tested antifungal ability by *in vitro* plate assays, focusing on the passaged lines after 10 cycles. We did not observe general phenotypic or genotypic changes in insect-passaged CHA0 in comparison to the ancestor, neither in insecticidal activity, nor in any other assessed phenotypic trait (Fig 2A-B,Fig 5, and S4.1-19). In addition to the insect lines, also all background lines preserved their insecticidal ability, and all lines — both insect and background lines — remained completely competent in plant root colonisation and plant protection. This suggests that the multi-host lifestyle of CHA0 was preserved during the serial passage experiment.

Evolutionary changes in bacterial lifestyles have been investigated in numerous serial passage experiments, focusing either on host switching [38,44,45] or on shifts in the nature of the bacteria–host interaction itself [34,36,39,46–51]. Studies examining bacterial adaptation to insect hosts through repeated cycles of feeding to insects and re-isolation on selective media have yielded contrasting outcomes, ranging from stable interactions to attenuated or increased virulence [36,46,48,50]. Two studies reported results comparable to ours. Schröder et al. [50] observed no change in the bacteria-host interaction following serial passaging of a human probiotic *Escherichia coli* strain within larvae of the insect host, and Mikonranta et al. [46] found that evolution in the absence of the host did not alter virulence of the opportunistic pathogen *Serratia marcescens* towards the insect.

In contrast, host adaptation experiments involving different *Pseudomonas* species, including CHA0, have reported either reduced virulence towards plant or nematode hosts [34,39] or enhanced colonisation capacity relative to the ancestral strain following passaging [38,51]. Collectively, these studies demonstrate that serial passaging frequently results in measurable adaptation to the imposed selection regime. However, no consistent trend emerges with respect to the direction of virulence evolution in insect hosts. Changes in virulence, whether attenuation or enhancement, appear to depend on a combination of biological factors, such as genetic architecture and host specificity [30], and experimental design parameters, including bottleneck size, duration of passaging, and selection criteria [52].

### Interpreting the lack of host-adaptation

In the following paragraphs, we address several aspects that may contribute to the lack of host adaptation in our study. First, our system exposed CHA0 to a complex and non-sterile environment whereas most previous serial passage studies were conducted in sterile or highly constrained environments, such as the plant vascular system. Under such conditions, bacteria may reduce investments in traits required for inter-microbial competition as competition is largely absent. Loss or downregulation of these competitive traits can have pleiotropic consequences, potentially reducing virulence or altering host interactions [39,53]. Indeed, even the presence of a single additional bacterial species or a simplified artificial microbiome has been shown to alter evolutionary trajectories in serial passage systems involving *Caenorhabditis elegans* and bacterial pathogens [33,47,49]. Repeating our experiment under fully sterile conditions could reveal whether constrains linked to direct competition affect niche specialisation. Likewise, serial passage of CHA0 in plant root systems with and without a resident microbiota may help disentangle trade-offs between host association and inter-microbial competition.

Second, relatively few studies have imposed explicit selection based on host health status during propagation [33,36,37,47]. In our experiment, we consistently selected moribund larvae, thereby directly selecting for virulence-associated traits. However, the evolutionary consequences of selecting diseased hosts, healthy hosts, or applying no host-based selection at all are likely to differ substantially [52]. This consideration may be particularly relevant in our system, where the outcome of the CHA0-insect interaction is strongly context-dependent [21,29]. For example, earlier work demonstrated that the infection outcome in *P. xylostella* depends on the initial bacterial dose: high doses led to host mortality, whereas lower doses allowed larval survival until adult emergence despite persistent colonisation [21]. Reducing the infection dose and selecting for healthy pupae could therefore have favoured a shift towards commensalism rather than maintained virulence, representing an alternative evolutionary trajectory worth exploring.

Third, our experimental design imposed multiple, potentially counteracting selection pressures that were absent in studies without regrowth steps [39,44,45,51,54]. In our system, bacteria experienced selection during growth on selective agar, survival on artificial food pellets, and infection of the insect host. On selective media, rapid growth and antibiotic resistance likely constituted dominant selective forces, potentially favouring fast-growing mutants. Such mutants commonly arise during prolonged culturing in nutrient-rich conditions [55,56]. We attempted to mitigate this by reducing nutrient concentrations to one-third strength, but selection for growth efficiency likely persisted. Subsequent exposure on food pellets introduced additional stresses, including desiccation and microbial competition, before ingestion by larvae. Inside the insect, bacteria encountered further barriers such as pH variation, oxygen stress, immune responses, and translocation into the haemolymph [15–20]. It is conceivable that only a small fraction of cells successfully established haemolymph colonisation [21], implying repeated bottlenecks and potential, random loss of beneficiary mutant strains. However, we did not detect strong genomic bottlenecks (Fig 6). At the same time, rapid fixation of several synonymous mutations within two cycles (Fig 3) does suggest strong selection or repeated reductions in effective population size that were insufficient to leave a clear genomic signature. Together, these observations indicate a complex interplay between selection and population dynamics, but do not provide unequivocal support for strong bottleneck-imposed population effects.

Although host adaptation and evolutionary trade-offs have been reported in other serial passage studies, including work with *P. protegens* CHA0, our results suggest that either intrinsic constraints such as a tight regulation of gene expression and the redundant use of factors in different environments stabilize the multi-host lifestyle or that the selection pressure in our system was insufficient to drive rapid specialisation within ten passages. The absence of marked phenotypic or genotypic shifts may therefore reflect evolutionary robustness rather than simply weak selection. Extending the number of passage cycles would clarify whether this stability persists under prolonged selection. Future experiments with other *Pseudomonas* species, particularly members of the *Pseudomonas chlororaphis* group that exhibit greater diversity in insect-associated traits [13,22], will be important to determine whether the preserved multi-host lifestyle observed here is a general feature of insecticidal pseudomonads or specific to CHA0.

### Different mechanisms might contribute to changes in some of the independent serial passage lines

While no general patterns of adaptation were observed after repeated passaging, several individual independent replicate lines exhibited phenotypic alterations. The increase in insect killing speed in some lines appears to be linked to specific mutations in the tested lines (Fig 5, S5.1), thus providing insights into the possible molecular mechanisms CHA0 might use to infect insect larvae or persist within the serial passage system.

Possible mechanisms underlying insecticidal activity may be identified through an investigation of the three lines Bkg1, Ins5, and Ins8, which displayed an increase in killing speed (Fig 2B,Fig 5). Ins8 showed a mutation in *choX*, encoding the periplasmic binding protein of a choline ABC transporter (Table 1). The degradation of phosphatidylcholine (PC) is a host-colonisation factor of *Pseudomonas aeruginosa* supporting replication in the respiratory tract [57,58]. The bacteria start PC degradation with the extracellular breakdown of PC into choline, glycerol, and fatty acids, which are subsequently taken up and metabolised within the bacterial cells [59]. Deletion mutants of different fatty acid degradation systems replicated less in the mice respiratory system compared to the wildtype [59] and choline uptake was speculated to be involved in *in vivo* survival [57]. Similarly, increasing the uptake of choline into the cell might be an adaptive strategy of CHA0 to unlock PC as an additional resource during larval infection. This hypothesis is supported by the finding that carbohydrate and amino acid transport and metabolism were the most enriched COG categories among genes carrying mutations (Fig 3A). Metabolic stress could thus be one of the challenges CHA0 encounters during insect colonisation. Additionally, the deletion of phospholipase C, responsible for extracellular degradation of PC, reduced insecticidal activity of CHA0 [12] and the expression of *choX* was upregulated during early infection of *P. xylostella* larvae [15]. Both findings support the hypothesis that PC degradation might contribute to the insecticidal activity of CHA0. The other two lines with an increase in killing speed, Bkg1 and Ins5, have a mutation in *lys1_2* encoding a saccharopine dehydrogenase or are not clearly associated with a specific mutation, respectively. Ins5 showed a tendency towards increased larval colonisation levels, which might result in the observed faster killing speed. However, this was not observed for Bkg1.

We found several lines with phenotypic changes in *in vitro* bacterial inhibition, swarming and growth characteristics, which might be connected to mutations in *PPRCHA0_1039* encoding a LTTR and in *PPRCHA0_2684* putatively encoding RstB (Fig 4D-G, Fig 5). Both coding regions were mutated in insect as well as background lines, indicating that they contributed to survival on the pellet and the selective medium rather than inside the insects.

Li et al. [39,60] and Song et al. [61] reported several independent mutations in the GacS/GacA two-component signal transduction system in evolution experiments with CHA0 or its close relative *P. protegens* Pf-5 which were associated with mutualistic and swarming traits, respectively. The Gac-Rsm network is a global regulatory system in *Pseudomonas* spp. which integrates environmental and cellular changes and induces lifestyle changes [26,27]. GacS/GacA deficient (Gac^-^) mutants appear spontaneously and reversibly during prolonged growth of *Pseudomonas* spp. on rich medium or in root systems [55,56,62,63]. GacS/GacA was not affected in our serial passage experiment except in one single colony (Table 1). There might be different reasons for the absence of mutations affecting GacS/GacA in our experiment. *gacS* or *gacA* deletion mutants show strongly reduced oral insecticidal activity [10–12,17] making Gac^-^ mutants unlikely to persist or be selected for in our serial passage system. Additionally, in contrast to the above-mentioned serial passage studies, our system was not sterile. Yan et al. [55] demonstrated that the accumulation of spontaneous Gac^-^ mutants was reduced in co-cultures with other bacteria indicating that the production of antimicrobial agents under the control of GacS/GacA are essential for competition.

By connecting phenotypes with genotypes, we gained initial insights into yet unknown mechanisms that enable CHA0 to survive in and colonise insects. The inclusion of a background control was critical, as it allowed us to disentangle mutations that promote insect colonisation and killing from those conferring advantages on the pellet or selective medium. Future experiments should aim to further characterise the mechanisms underlying the observed phenotypes. For example, testing knock-out mutants of the PC degradation pathway for changes in insecticidal activity could reveal its contribution to CHA0’s multifactorial insecticidal capacity. Similarly, functional validation of the putative targets of *PPRCHA0_1039* would help to understand its involvement in the background adaptation process.

## Conclusion

The evolutionary trajectories of host-relationships that bacteria undergo during serial passage experiments vary extensively and depend on a multitude of factors. In our study, *Pseudomonas protegens* CHA0 remained largely unchanged after serial passaging through insect hosts. It neither lost nor gained insecticidal or plant-protective traits, and therefore preserved its insect- and plant-associated, multi-host lifestyle. This evolutionary stability across complex and fluctuating conditions — such as those encountered in agricultural ecosystems — supports the potential of CHA0 as a biocontrol agent in the field and makes CHA0 a promising candidate for reliable biological control of plant diseases and insect pests.

## Materials and methods

### Serial passage experiment

Serial passage of *Pseudomonas protegens* CHA0 was done with ten independent replicate lines to allow independent evolution in larvae of *Plutella xylostella* (Ins4-13) and three independent replicate background lines without larvae present (Bkg1-3) (Fig 1). To start the lines, *P. protegens* CHA0-gfp (see Table 2 for all microorganisms used in this paper) was grown on one-third strength King’s B medium (20 g proteose peptone, 8.4 ml 87% glycerol, 1.5 g MgSO_4_ x 7H_2_O, 1.5 g K_2_HPO_4_ x 3H_2_O dissolved in 1 l ddH_2_O) [64] agar supplemented with 13 µg/ml chloramphenicol, 100 µg/ml cycloheximide, and 10 µg/ml gentamycin (1/3 strength KB^++G^) for two days at 24°C to recover the cells from storage at -80°C in 43% glycerol. Cell material was scraped from the plate and re-suspended in 2 ml 0.9% NaCl solution (saline). 10-fold serial dilutions in saline were plated on 1/3 strength KB^++G^ agar and incubated for 4 days at 18°C to achieve sufficiently high cell numbers to start with. All cell material was scraped from the agar plate, on which single colonies were just visible (on average 4*10^4^ colonies, see Fig S1.1C for detailed numbers of pooled colonies), re-suspended in 50 ml saline, washed once (one volume saline, centrifugation for 10 min at 4°C and 2700 rpm (Allegra X12R Centrifuge, Rotor: SX4750, Beckman Coulter Life Sciences)) and finally re-suspended in 5-10 ml saline. For DNA extraction, aliquots of 2-4 ml cell suspension were spun down (11’000x g, 2 min), supernatant was removed, and the pellets were frozen at -80°C. For long-term storage, aliquots of 1 ml cell suspension were mixed 1:1 with 87% glycerol and stored at -80°C. For inoculation, the cell density was adjusted to an OD_600_ of 0.2 (= 3*10^8^ CFUs/mL) and the same ancestor inoculum was used to start all lines. *P. xylostella* rearing, infection and monitoring were performed as described by Flury et al. [17]. Briefly, eggs obtained from Syngenta Crop Protection AG (Stein, Switzerland) were reared on Beet Armyworm Diet with vitamins but without chlortetracycline (Frontier Agricultural Sciences, Newark, DE, USA; Ref: F9220B) in a climate chamber (60% relative humidity; day: 16 h, 25°C, 12 kLux; night: 8 h, 20°C) and starved for 3-4 h prior to infection. Pellets of artificial food were placed on wet filter papers in 128-well insect trays. For the insect lines, pellets were inoculated with 3*10^6^ CFUs and one second instar larvae was added per well. For the background lines, pellets were inoculated exactly the same but no larvae were added. Ten insect lines with 30 larvae each and three background lines with 12 pellets each were run in parallel. After 3 days, five moribund larvae per insect line were surface-disinfected by subsequently dipping larvae for 20 s in each 70% EtOH, 0.05% SDS, 70% EtOH and ddH_2_O. Lavae were dried on filter paper, pooled and homogenised in 100 µl saline using a 2 ml Eppendorf tube, one heat-sterilised glass bead per tube (ø 5 mm) and the MM300 TissueLyser (Retsch GmBH, Haan, Germany) for 2 x 45 s at 30/s. Five pellets per background line were pooled and homogenised without surface-disinfection as described. Homogenates were diluted with 400 µl saline. 10-fold serial dilutions in saline were plated on 1/3 strength KB^++G^ agar, incubated for 4 days at 18°C (see Fig S1.1A for detailed numbers of recovered bacteria). Colonies were scratched from the plates for re-inoculating pellets of the next cycle and cell material was prepared for DNA extraction and long-term storage as described above.

After the last cycle, several aliquots of long-term stocks were additionally prepared for each line for phenotyping. Furthermore, twelve single colonies per line were picked from the 1/3 strength KB^++G^ agar plates and grown overnight in lysogeny broth (LB) at 24°C and 180 rpm. Cultures were mixed with glycerol and long-term stored as described above. These long-term stored single colonies were re-grown a second time overnight in LB before cultures were spun down as described above for DNA extraction.

Re-isolation of CHA0 from the background lines failed sometimes. When this happened, the glycerol stock of the last cycle was re-grown for 2 days on 1/3 strength KB^++G^ agar plates at 24°C and used as inoculum for the next round. This happened with line Bkg1 in cycle 1 and cycle 5 and with Bkg3 in cycle 5. To obtain a similar number of cycles as with the other lines, Bkg1 was further cycled until cycle 9 (hereafter referred to as cycle 10) and Bkg3 until cycle 10.

An approximation of the generations that CHA0 underwent on the food pellet or within the larvae was calculated using the formula below. For the background lines, the number of generations was estimated based on the size of the inoculated population (N_0_ = 3*10^6^ cells) and the population size measured at the extraction time point. For the insect lines, a slight bottleneck during the colonisation of the larvae was assumed (N_0_ = 10^5^ cells) which is consistent with the results of our bottleneck experiment and the colonisation densities reported by Vesga et al. [15] 24 hours after insect infection.

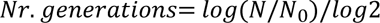

### Genotyping

#### Whole genome resequencing

The CHA0 populations of each independent replicate line from cycles 1, 2, 4, 6, 8, and 10 as well as 12 single colonies picked per line from cycle 10 were subjected to whole genome re-sequencing. Genomic DNA was isolated using the QIAamp DNA Mini Kit (Qiagen, Venlo, The Netherlands; Ref: 51306) and paired-end DNA libraries for Illumina sequencing were prepared using the Illumina DNA Prep kit (Illumina, San Diego, CA, USA; Ref: 20060059) according to the Hackflex protocol by Gaio et al. [67] (see Supplementary Information S1). Paired-end libraries were pooled accordingly and sequenced on the Illumina NovaSeq S1 2 x 150 platform at the Utrecht Sequencing Facility USeq.

#### Variant calling and enrichment analysis

Libraries were quality checked with MultiQC (version: 1.12) [68]. The most recent reference genome of CHA0 (parent strain of CHA0-gfp) available on NCBI (NCBI, LS999205.1) does not contain the miniTN7-tag of CHA0-gfp, therefore this sequence was reconstructed from the ancestral samples by *de novo* assembly using spades (version: 3.15.4) [69] and added to the available reference genome. The updated reference genome was used for variant calling with SNIPPY (version: 4.6.0) [70] which finds single nucleotide polymorphisms or small insertions and deletions and provides the affected coding regions if possible. SNIPPY was adjusted to identify mutations at intermediate allele frequencies by relaxing the default settings that retains fixed mutations. The ancestral CHA0 samples were used as a control to exclude detected mutations resulting from pre-existing mutations in the used long-term stock or from misalignments by SNIPPY (see Table ST1.2). Additionally, six mutations were removed as they had low read numbers and were randomly popping up in more than 3 lines without any clear patterns, mostly in non-coding sequences (see Table ST1.2). This allowed us to obtain the relative frequency of each mutation in the populations and the presence of mutations in the single colonies. A subset of the affected coding regions was subjected to literature research, an InterPro Scan [71] to find affected subdomains and 3D protein structures were modelled using the AlphaFold Server [72]. Coding regions hit by population-level mutations were used for COG and KEGG pathway enrichment analysis, for details see Supplementary Information S1. Population-level reads not mapping to the CHA0 reference genome were, where possible, assigned to other microbial taxonomic labels to detect potential contaminants accumulating during the cycles, for details see Supplementary Information S1.

### Phenotyping

The following bioassays were performed for phenotyping: *P. xylostella* feeding, *Pythium ultimum*-cucumber, fungal and bacterial inhibition, swarming, biofilm formation and *in vitro* growth assay. For details of the bioassays and the isolation of the *P. xylostella* gut isolates *Arthrobacter* sp. Px40 and *Microbacterium* sp. Px99 see Supplementary Information S1. Each line after cycle 10 and a subset of single colonies were tested in at least two independent experiments per assay. Relative values were calculated per experiment to minimise variations between experiments using the equation below and pooled from all experiments per assay for statistical analysis in R-studio (version: 2023.06.0). Relative values were compared using a linear mixed effect model (package: lme4 [73]) followed by post-hoc pairwise comparisons of estimated marginal means (package: emmeans) adjusting the *p*-value with the Tukey method for multiple testing. First, the insect and background passage treatments were compared to the ancestor, then, each individual independent replicate line was compared to the ancestor.

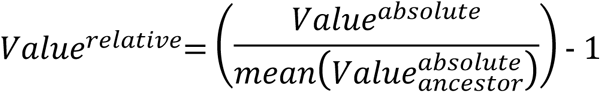

Larval survival in the *P. xylostella* feeding assays was analysed over relative time starting with the inoculation and ending when 95-100% of the larvae were dead. Survival curves were compared using the cox hazards model (package: coxme [74]) followed by post-hoc pairwise comparisons of estimated marginal means (package: emmeans [75]) adjusting the *p*-value with the Tukey method for multiple testing. Alternative allele frequencies of all genomic regions hit by a mutation in the populations at cycle 10 were correlated with relative values of the respective phenotypes in R-studio (package: psych [76]). *P*-values were adjusted using the default of the package for multiple testing.

### Bottleneck experiment

Inoculums (OD_600_ = 0.02) of *P. protegens* CHA0-gfp and *P. protegens* CHA0-mche were prepared as described above in Serial passage experiment and mixed in the following ratios: 1:1, 1:10, 1:10^2^, 1:10^3^,1:10^4^. The ratios in the inoculum were verified by 10-fold serial dilutions in saline and spotting on KB^++G^ agar. Pellets of artificial food were inoculated and to one half of them *P. xylostella* larvae were added. Larval as well as pellet colonisation by each, CHA0-gfp and CHA0-mche, from six replicates was determined after 30 h as described in Serial passage experiment. The two strains were differentiated based on their emitted fluorescence by counting colonies with a Leica M205 FCA fluorescence stereomicroscope (Leica, Wetzlar, Germany). Data of the two independent experiments (Exp. 1: treatments: 1:1 to 1:10^3^, Exp. 2: treatments: 1:10 to 1:10^4^) were pooled for analysis in R-studio (version: 2023.06.0). CFUs were log_10_-transformed to calculate ratios. Points below the detection limit (5 CFUs/larva or pellet) were set to 2.5 CFUs/larva or pellet. The absolute difference of the larva or pellet ratios from the inoculum ratio (Deviation) was calculated using the equation below and compared using a general linear mixed model adjusting for gamma distribution followed by post-hoc pairwise comparisons of estimated marginal means (package: emmeans) adjusting the *p*-value with the Tukey method for multiple testing.

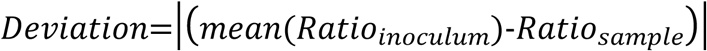

## Acknowledgements

We would like to thank Sabrina Müller, Fortesa Rama, Silja Müller, Lina Jäger, Ella Staub and Stojanka Mitrovic for their support with experimental work. We kindly thank Juan J. Sanchez-Gil (University of Utrecht) for support with the swarming assay and Petra Reiniger, Oliver Kindler, and their team (Syngenta) for steadily delivering *P. xylostella* eggs. Part of the data produced and analysed in this paper were generated in collaboration with the Genetic Diversity Centre (GDC), ETH Zurich. We acknowledge the Utrecht Sequencing Facility (USEQ) for providing sequencing service and data. USEQ is subsidised by the University Medical Center Utrecht and The Netherlands X-omics Initiative (NWO project 184.034.019).

## Supporting information

S1. Supplementary Methods and Figures.

ST1.1. Additional information on all mutations.

ST1.2. Additional information on all ancestral and random mutations.

